# Dissecting phenotypic transitions in metastatic disease via photoconversion-based isolation

**DOI:** 10.1101/2020.10.05.327213

**Authors:** Yogev Sela, Jinyang Li, Paola Kuri, Allyson Merrell, Ning Li, Chris Lengner, Pantelis Rompolas, Ben Z. Stanger

## Abstract

Cancer patients presenting with surgically resectable disease often harbor occult metastases, a potential source of relapse that is targetable only through systemic therapy. Studies of this occult fraction have been limited by a lack of tools with which to isolate discrete cells based on spatial grounds. We developed PIC-IT, photoconversion-based isolation technique allowing efficient recovery of cell clusters of any size including solitary disseminated tumor cells (DTCs), which are largely inaccessible otherwise. In a murine pancreatic cancer model, transcriptional profiling of spontaneously arising DTCs revealed phenotypic heterogeneity, functionally reduced propensity to proliferate and enrichment for inflammatory-response phenotype associated with NF-κB /AP-1 signaling. Pharmacological inhibition of NF-κB depleted DTCs but had no effect on macrometastases, suggesting DTCs are particularly dependent on this pathway. PIC-IT enables systematic investigation of the earliest stages of metastatic colonization. Moreover, this new technique can be applied to other biological systems in which isolation and characterization of spatially distinct cell populations is not currently feasible.

## Introduction

Metastasis is the primary cause of cancer-associated mortality and remains a significant therapeutic challenge. A considerable fraction of metastasis is occult (Haeno et al., 2012; Vanharanta & Massagué, 2013), and thus serves a potential source for residual disease and recurrence (María Soledad Sosa, Bragado, & Aguirre-Ghiso, 2014; Tohme, Simmons, & Tsung, 2017). While fluorescent reporters allow detection of metastatic colonies in pre-clinical models (Aiello et al., 2016; Fluegen et al., 2017), an inability to recover pure micron-scale colonies has precluded systematic phenotypic analysis of early stage metastases. Several methods have been developed to isolate cells from specific compartments *in vivo*, laser-capture microdissection (LCM) representing the most widely used platform (Basnet et al., 2019; Espina et al., 2006; Lovatt et al., 2014; Tang et al., 2009). However, the use of these methods is restricted to compartments of particular size and only for specific applications due to several limitations, including: (i) contamination by undesired cells within the capture field; (ii) low throughput; (iii) a need for expensive and temperamental hardware and (iv) loss of cell viability. (v) Lack of control over population composition. Spatial transcriptomics methods continue to improve and can distinguish local patterns across large tumor regions (Moncada et al., 2020), but resolution limits and low representation of small metastatic colonies in the tissue hinders their isolation. Photoactivable and photoconvertible proteins provide a promising alternative for targeted cell isolation (Medaglia et al., 2017; Nicenboim et al., 2015). However, existing systems require specialized equipment and extended handling times, thus limiting scalability and precluding effective acquisition of rare or sporadic cells from multiple compartments-such as DTCs. Here, we report “PIC-IT” (“Photomark Isolate Cells If Tiny”) which enables unbiased and efficient isolation of size-defined metastatic colonies from live tissues through photoconversionbased marking (Figure 1A).

**Figure 1.**
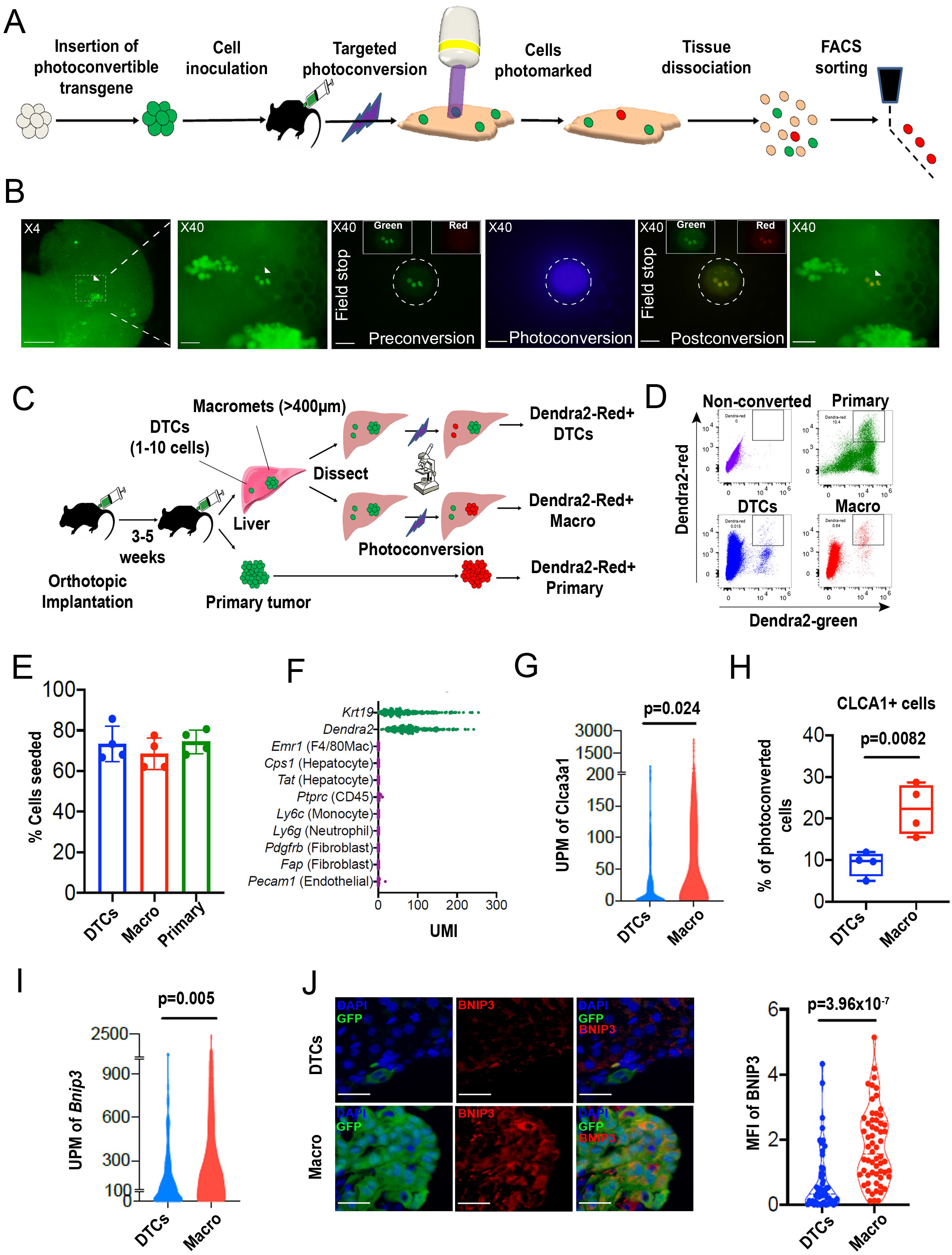
PIC-IT allows isolation of size-defined DTCs and macrometastasis from live animals. **(A)** Schematic of the PIC-IT protocol. **(B)** Photoconversion of Dendra2 in DTCs spontaneously arising in the liver of a pancreatic cancer tumor model. Left to right: Microscopic views acquired through a 4X objective with region of interest containing 3-cells cluster destined for photoconversion located in the marked area (Arrows denote the 3-cells of interest, Scale Bar=250μm). Focus on the marked area with 40X objective (Scale Bar=25μm). Field stop-confined FOV within the 40X objective region, encircling the targeted 3-cell focus prior to photoconversion (“Preconversion”). In boxes, images representing individual green and red channels for the field-stop FOV. Field-stop image of a photoconversion session (violet light exposure, “Photoconversion”). Field-stop confined FOV of an X40 objective following photoconversion (“Postconversion”). The X40 region with a full FOV demonstrating specificity of photoconversion to field-stop confined area. **(C)** Experimental scheme utilized for photoconversion-based isolation of metastatic cells from liver fragments. Orthotopic tumors derived from Dendra2-expressing cells produce liver metastasis. In different fragments, DTCs or macrometastasis are photoconverted. Dissociated tissue from each group are pooled and sorted for subsequent applications. **(D)** Flow cytometry charts showing expression of Dendra2-green (X-axis) and Dendra2-red (Y-axis) in converted tumor cells and non-converted controls. **(E)** Seeding efficiency of photoconverted tumor cells, calculated as the fraction of cells counted in tissue culture plates 24h post-isolation relative to the number sorted by FACS (n=4). **(F)** UMI profiles indicate expression of the markers *Krt19* and *Dendra2* in the sorted cells but no expression of other lineage markers (N=361). **(G,H)** High prevalence of RNA-seq transcripts of *CLCA3A1* **(G)** (N_DTCs_=94, N_macro_=111) and the corresponding CLCA1 protein expression **(H)** determined by flow cytometry on macrometastasis-derived vs. DTCs (n=4). **(I,J)** Concordance between elevated transcription (UMI per million) of BNIP3 in RNA-seq **(I)** and **(J)** BNIP3 MFIs in (Left panel) immunostained DTCs and macrometastasis populating the livers of 6419c5-YFP tumor bearing mice (N_DTCs_=53, N_macro_=60). On the right, representative images of DTCs and macrometastasis immunostained for BNIP3 (Red), YFP (green) and DAPI (Blue). Scale bar=50μM. Bars represent mean ± SEM in all graphs. **, *p*<0.01; ***, *p*<0.001 by p-values were calculated by unpaired two-tailed Student’s t-test.

## Results

### Photoconversion-based isolation of size-specific metastatic colonies

Photoconvertible proteins comprise a class of fluorescent proteins whose emission switches from green to red upon exposure to blue light. For these studies, we used a version of the photoconvertible protein Dendra2 fused to H2B (H2B-Dendra2). As shown in Figure1-figure Supplement 1A exposure of H2B-Dendra2-expressing pancreatic tumor cells (5074 cell line) to a mercury lamp-generated violet light (400-450 nm) for 30 seconds efficiently marks cells. Next, we engineered a highly metastatic pancreatic cancer cell line (6419c5) to express H2B-Dendra2. Use of a clonal cell line was chosen to minimize genetic-based contribution to metastatic heterogeneity (Hunter, Amin, Deasy, Ha, & Wakefield, 2018). To generate metastasis, 10^4^ 6419c5-H2B-Dendra2 cells were injected into the pancreas of immune competent C57BL/6 mice and tumors were allowed to grow for 3-5 weeks, at which point livers were examined under a standard widefield microscope.

Imaging revealed metastatic foci of various sizes, ranging from isolated disseminated tumor cells (DTCs) to micrometastases and larger macrometastasis (Figure 1B, left). Importantly, strong nuclear fluorescence in all tumor cells enabled easy detection of metastatic foci of a particular size even with a low-magnification 4X objective, allowing for rapid identification of regions of interest (ROIs). To test whether region-specific photoconversion could be achieved using a widefield microscope in unsectioned, intact tissue specimens, we confined the field of view (FOV) using the field stop element (Figure 1B, “Preconversion”) and exposed the cells to 400-450 nm light (Figure 1B, “Photoconversion”). Exposure for <5 seconds (40X objective, N.A.=0.75) led to a robust induction of red fluorescence in all cancer cells residing specifically within the boundaries of the limited FOV (Figure 1B, “Postconversion”). In a different FOV, using a 10X objective, we rapidly photoconverted (15 seconds, N.A=0.4) hundreds of cancer cells within a macrometastasis (Figure1-figure supplement 1B). Spatial precision of labeling was achieved even when a low magnification objective was used, with a rapid drop-off of photoconversion outside the FOV boundary (Figure1-figure supplement 1C). These results demonstrate that both small and large metastatic lesions can be efficiently photoconverted using a wide-field microscope by adjusting the FOV, exposure time, and numerical aperture.

We next tested whether we could recover photoconversion-marked metastatic cells for flow cytometry-based applications. To maximize accessibility to metastatic foci, we dissected the liver into millimeter-sized fragments and photoconverted cells within each fragment separately (Figure 1C). Consequently, each liver fragment was designated for photoconversion of DTCs (Figure 1C, top) or macrometastases (Figure 1C, middle). In parallel, cells in the primary tumor were also photoconverted (Figure 1C, bottom). This strategy allowed samples to remain chilled throughout the procedure, with exposure to ambient temperatures only during the photoconversion process (<6 minutes). DTCs were defined as collections of 10 or fewer metastatic cells, with the majority being 1 or 2 cells (Figure1-figure supplement 2A) while macrometastases were defined as lesions of >400μm in diameter. Following flow sorting, photoconverted cells derived from the different groups showed strong, comparable increase of ~30-fold in red fluorescence intensity (Figure 1D and Figure1-figure supplement 2B). Although red fluorescence increased with greater light exposure times, 5 sec was sufficient to detect the photoconverted cells (Figure1-figure supplement 2C). These results indicate that tissue resident H2B-Dendra2-expressing tumor cells can be readily detected by flow cytometry following brief photoconversion and tissue dissociation.

Because low throughput has been one of the key limitations impeding studies of DTCs, we next sought to quantify the throughput of our method. To this end, we photoconverted 200-400 DTCs in each of 16 different isolation sessions (~2.5 hours per session) and determined the yield of photoconverted tumor cells by flow cytometry. Across these sessions, recovery of photoconverted DTCs averaged at 38% (Figure1-figure supplement 2D), suggesting that 80-170 cells can be acquired per hour using this experimental setup (Figure1-figure supplement 2E). Notably, we typically used less than 30% of the liver in our marking sessions. Cell yield could potentially be improved with longer sessions, as there was minimal decay in the red signal up to 7 hours post photoconversion (Figure1-figure supplement 2F). Cells remained viable after flow sorting, with a ~70% seeding efficiency measured 24 h post plating (Figure 1E). Importantly, cells sorted following photoconversion showed no evidence for DNA damage (Figure1-fig supplement S3A) or impairment of colony formation (Figure1-figure supplement 3B). Overall, these results demonstrate robust acquisition of hundreds of live cells from size-defined metastatic foci within a practical timeframe.

Single-cell RNA-seq of 361 metastatic and primary tumor cells via Cel-seq2 (Hashimshony et al., 2016) revealed expression of dendra2 and *Krt19* but not markers of hepatocytes, endothelial cells, immune cells and fibroblasts (Figure 1F), confirming the purity of the isolated cell populations. To further validate this RNA-seq expression analysis, we identified two genes – *Clca3a1 and Bnip3* – predicted to be more highly expressed in macrometastases than DTCs and confirmed their enrichment by flow cytometry and immunostaining in mice injected with 6419c5-YFP cells (Figure 1G-J). Taken together, our findings suggest that PIC-IT enables efficient, pure isolation of size-defined metastatic colonies and enables discovery of genes whose expression differs in the two populations.

### PIC-IT identifies enrichment for a pro-inflammatory phenotype in liver DTCs

In various experimental settings, DTCs have been associated with physiological adaptations related to their dissemination (Fluegen et al., 2017) and surrounding microenvironment (Linde, Fluegen, & Aguirre-Ghiso, 2016; Milette, Sicklick, Lowy, & Brodt, 2017). However, a systematic analysis of the phenotypic diversity of spontaneously arising DTCs is currently lacking. We thus examined the transcriptional profiles of PIC-IT-extracted DTCs and macrometases by Seurat clustering analysis (Stuart et al., 2019) and visualized by UMAP dimensional reduction (McInnes, Healy, Saul, & Großberger, 2018). At a global level, the distribution of DTC and macrometastastic transcriptomes broadly overlapped (Figure 2A). UMAP further identified 3 clusters in DTCs and macrometastasis, annotated as proliferative, inflammatory or hypoxic/stressed suggesting that DTCs are phenotypically diverse (Figure 2B and figure supplement 1A). Notably, the inflammatory cluster, defined by elevated IFN-γ response and complement activity (Figure2B and Figure2-figure supplement 1B), was enriched by 2-fold in DTCs comparing to macrometastasis (30% vs. 16%). Next, we carried a differential gene analysis using EdgeR (*17*) – comparing the transcriptomes of 94 DTCs to 111 macrometastatic cells. This – revealed 500 genes that were more highly expressed in DTCs and 366 genes that were more highly expressed in macrometastases (P<0.05, Figure 2B). Functional annotation of the DTC-associated genes revealed a strong signal for TNF-related NF-κB signaling and a prominent increase in inflammatory cytokine secretion (Figure 2C). Likewise, an IFN-γ response signature was enriched in DTCs, falling in line with the pattern of enriched inflammatory gene expression. Furthermore, transcription factor motif analysis (HOMER) revealed enrichment of pro-inflammatory DNA regulatory elements in DTCs (Figure 2D). In addition to the known inflammation-related transcription factors NF-κB, JUN/AP-1, and STAT, motif analysis also revealed enrichment of binding sites for MAFK which has-recently been demonstrated to enhance NF-κB transcriptional response (Hwang et al., 2013). In line with this enrichment of TNF-related NF-κB signaling, 29 genes associated with the Hallmark “TNFa-signaling-via-NF-κB” gene set were expressed at significantly higher levels in DTCs compared to macrometastastic cells (Figure 2E and Figure2-figure supplement 1A). Taken together, transcriptional profiling suggests that pancreatic DTCs in the liver manifest a strong pro-inflammatory phenotype associated with increase in NF-κB and AP-1 signaling.

**Figure 2.**
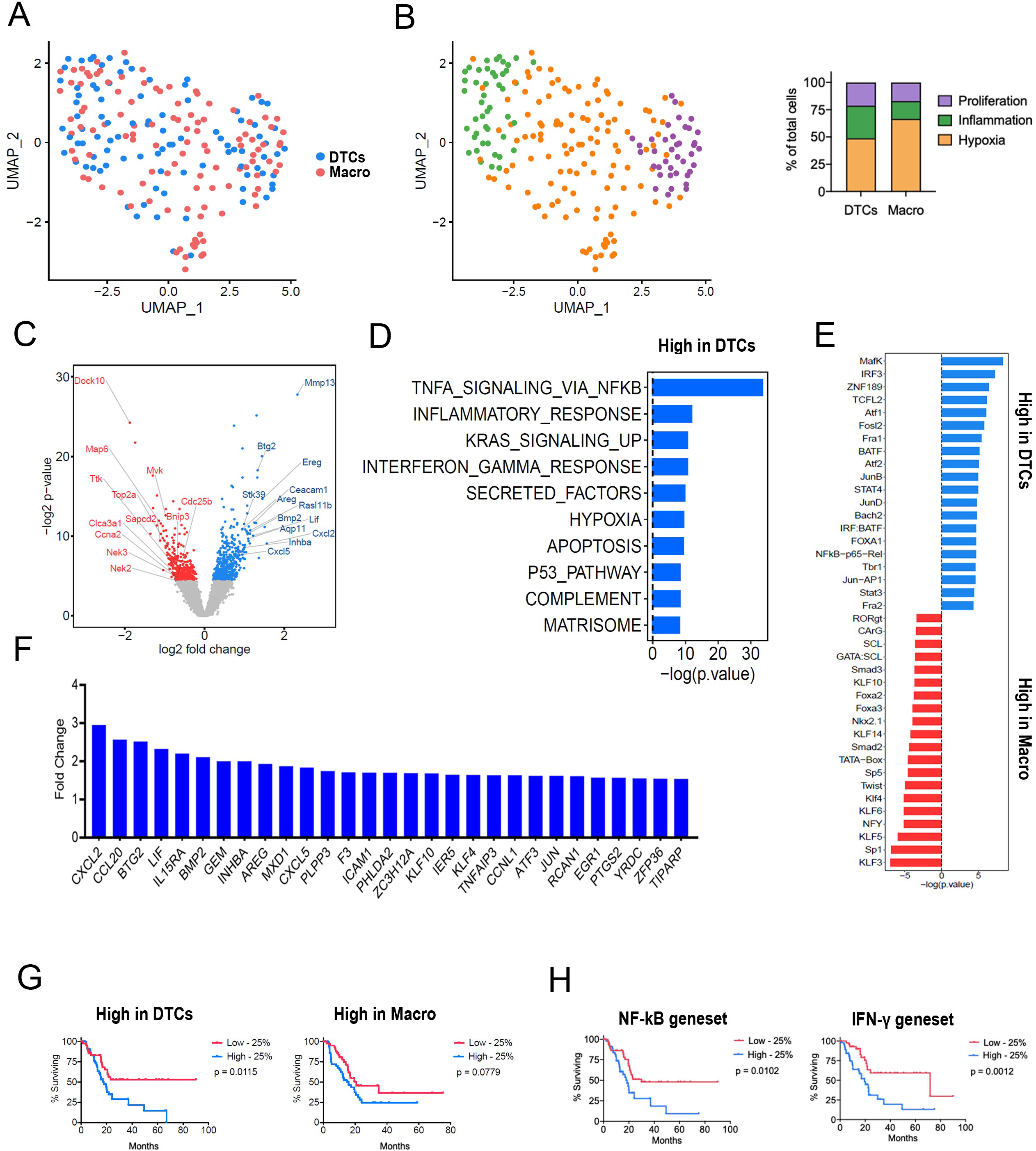
Phenotypic transitions in pancreatic tumor cells colonizing a mouse liver. **(A)** Transcriptome distribution visualized by UMAP (**B)** Distribution of metastatic cells for the different clusters functionally annotated by GSEA msGDMIB (p<0.05, 200 highest fold-change genes). On the right, proportions of clusters in DTCs and macrometastasis. **(C)** Volcano plot of differentially expressed genes in DTCs and large metastasis derived from differential gene analysis (EdgeR). **(D)** Functional annotation of genes upregulated in macrometastasis. GSEA msGDMIB for Hallmark and canonical pathways (p<0.05, 200 highest fold-change genes). **(E)** Motif analysis of potential transcriptional regulator using HOMER with −1000 bp to +300 bp as scanning region and p<0.05, 200 highest fold-change genes as input (showing knownResults with p<0.01) **(F)** Fold change increase (DTCs vs. Macrometastasis) in differentially expressed genes (p<0.05) associated with Hallmark TNFa-signaling-via-NF-κB signaling category derived from GSEA msGDMIB analysis. **(G)** Kaplan-meier curves showing overall survival of PDAC patients with tumors enriched for DTCs and macrometastasis signatures (p<0.05, 200 highest fold-change genes) and **(H)** curves comparing PDAC patients signatures to inflammatory-response signatures (N=177, Lowest vs. Highest quartiles, log-rank test).

To explore how the inflammatory profile of murine DTCs may relate to human pancreatic cancer, we created DTCs and macrometastasis gene scores from the respective gene signatures and applied them to TCGA data. Interestingly, our analysis that enrichment for DTCs-high genes but not macrometastasis-high genes predicted a poor overall survival in PDAC patients (Figure 3F). Specifically, enrichment, signatures associated with NF-κB transcriptional response and IFN-y-response pin DTCs individually poor prognosis (Figure 3G). These results suggest that concurring metastatic colonies may have stage-dependent impact on human disease and nominate contribution for the inflammatory response phenotype in prognosis.

**Figure 3.**
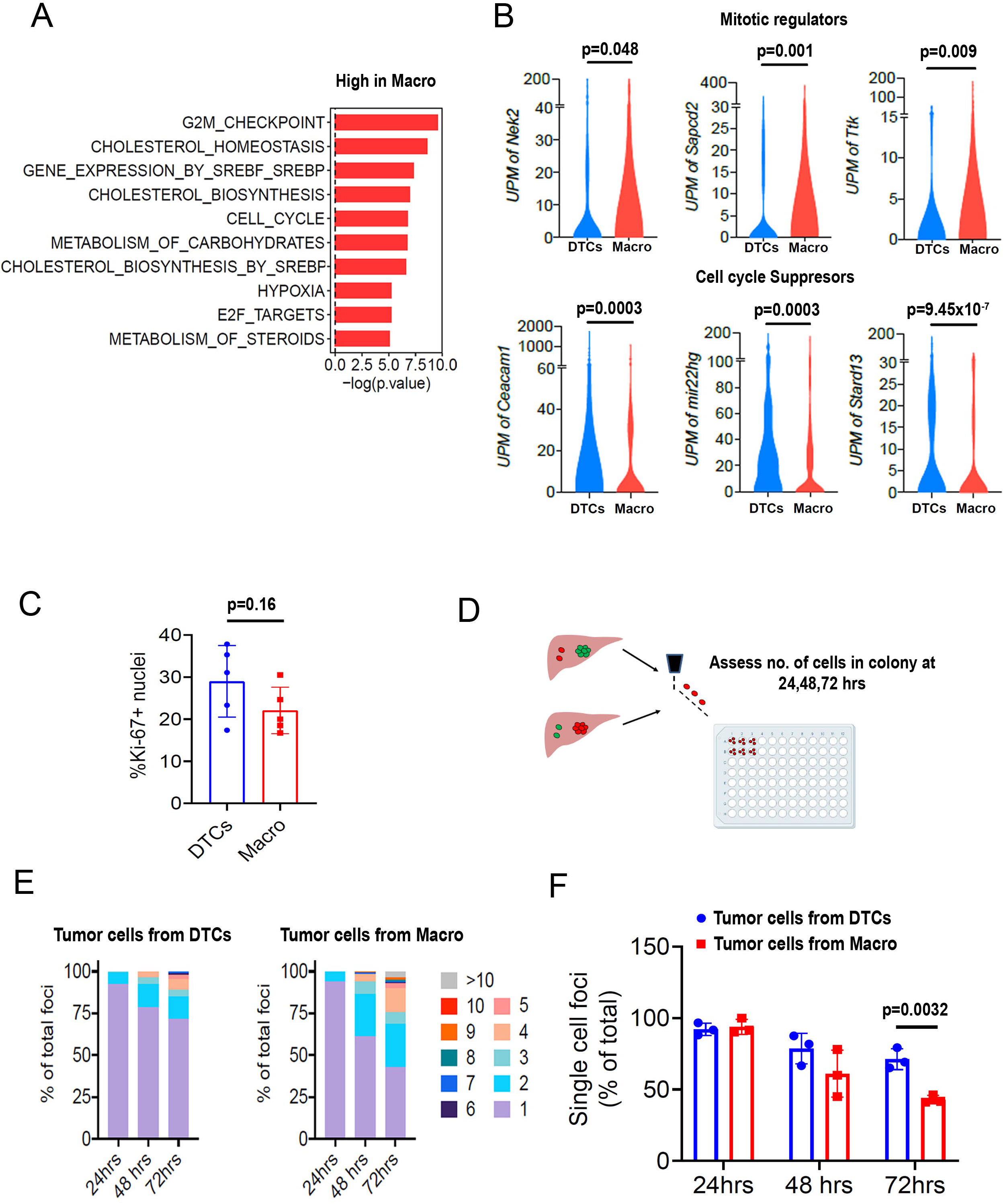
DTCs exhibit a hypo-proliferative phenotype. **(A)** Functional annotation of genes downregulated in DTCs/higher in Macrometastasis. GSEA msGDMIB for Hallmark and canonical pathways (p<0.05, 200 highest fold-change genes). **(B)** Abundance of transcripts in RNA-seq for genes associated with cell cycle arrest (Top) and positive regulation of mitosis (Bottom). (N_DTCs_=94, N_macro_=111). **(C)** Fraction of Ki-67 positive cells in DTCs and macrometastasis determined by immunostaining (n=5 samples per group). **(D)** Schematics for *ex vivo* proliferation assay. DTCs and macrometastasis are extracted via PIC-IT, seeded at 45 cells/96well and colony growth is monitored via microscopy over 72hrs. Image was made by Biorender. **(E)** Colony size distribution of metastasis-derived cells in the first 72h after isolation. Left panel, colonies derived from DTCs. Right panel, colonies derived from macrometastasis (n=3 independent experiments). **(F)** Fraction of isolated metastatic cells residing as solitary cells at 24, 48 and 72hrs post isolation (n=3 independent experiments). Bars represent mean ± SEM in all graphs. p-values were calculated by unpaired two-tailed Student’s t-test

### DTCs exhibit reduced propensity to proliferate

We next turned our attention to the genes with low expression in DTCs relative to macrometastases. Intriguingly, functional annotation indicated enrichment for cell cycle and cholesterol biosynthesis pathways in the macrometastases (Figure 3A), with elevated expression of a number of mitosis-regulators such as *Nek2* (Fry, O’Regan, Sabir, & Bayliss, 2012)*, Sapcd2* (X. Xu et al., 2007) and *Ttk* (Dominguez-Brauer et al., 2015) (Figure 3B). Consistent with this, DTCs exhibited higher expression of negative regulators of cell cycle progression including *Ceacam1* (Sappino et al., 2012), *Stard13* (Jaafar, Chamseddine, & El-Sibai, 2020) and *Mir22hg* (J. Xu et al., 2020) (Figure 3B), predicting an overall hypo-proliferative state for DTCs. Surprisingly, however, Ki-67 immunostaining revealed no measurable difference in the percentage of positive cells in DTCs vs. macrometastases (Figure 3C), consistent with our previously reported findings (Aiello et al., 2016).

We hypothesized cells in larger metastatic lesions may be inherently “poised” to proliferate more readily than those in smaller metastatic lesions, a difference that might not be reflected in elevated proliferation rates *in situ*. To test this, we exploited PIC-IT’s capacity to isolate live cells and functionally measured the proliferative potential of the different metastatic groups. DTCs and macrometastasis-derived cells were extracted from mice bearing 6419c5-H2B-Dendra2 tumors, seeded at a density of 45 cells/well, and monitored daily for colony growth (Figure 3D). While cells derived from DTCs and macrometastases expanded to a similar extent after 24h, colonies derived from macrometastases were larger at 48h and 72h (Figure 3E-F). These results suggest that despite similar rates of proliferation *in vivo*, macrometastatic cells may be primed for cell division (or, stated otherwise, that DTCs may be primed for cell cycle arrest).

### DTCs require NF-κB /AP-1 signaling

We have previously reported that solitary DTCs exhibit resistance to standard-of-care chemotherapy (gemcitabine + nab-paclitaxel) in autochthonous KPCY mice (Aiello et al., 2016). Consequently, we sought to identify vulnerabilities that might be specific to DTCs. As our profiling experiments indicated enrichment of a pro-survival inflammatory response in DTCs, we hypothesized that this pathway – driven by NF-κB and/or AP-1 – might be important for the persistence of these cells *in vivo*. First, we performed NF-κB-p65 staining in the 6419c5 implanted cell line model and the KPCY autochthonous model. These two models confirmed that DTCs exhibit higher levels of nuclear NF-κB-p65 compared to macrometastases (Figure. 4A-B). In addition, and as suggested from our motif analysis, phospho-c-Jun-positive cells were more prevalent in DTCs than macrometastases in both models (Figure4-figure supplement 1). These findings suggest that the enhanced inflammatory signatures identified through PIC-IT-enabled RNA sequencing reflect true signaling differences in small and large metastases.

To functionally test whether this pro-inflammatory response is required for DTC viability, we tested the potency of Triptolide (TPL), a potent anti-inflammatory agent that inhibits both pathways (Yuan et al., 2019). We inoculated 6419c5-YFP cells into the pancreas and allowed implants to grow for 21 days (Figure 4C), a time point at which >95% of mice develop macrometastasis (data not shown). Animals were then randomized to receive either Triptolide (0.2mg/kg) or vehicle by intraperitoneal injection (7d) followed by quantification of DTCs and macrometastases by immunohistology (Figure 4C). Although the total liver area occupied by metastases (dominated by macrometastases) was similar in the two treatment groups (Figure 4D), Triptolide treatment led to a significant reduction in the relative frequency of DTCs (Figure 4E) and the ratio of DTCs or single metastatic cells to macrometastases (Figure 4F,G). These results suggest that DTCs are more susceptible to inhibition of inflammatory signaling than macrometastasis.

**Figure 4.**
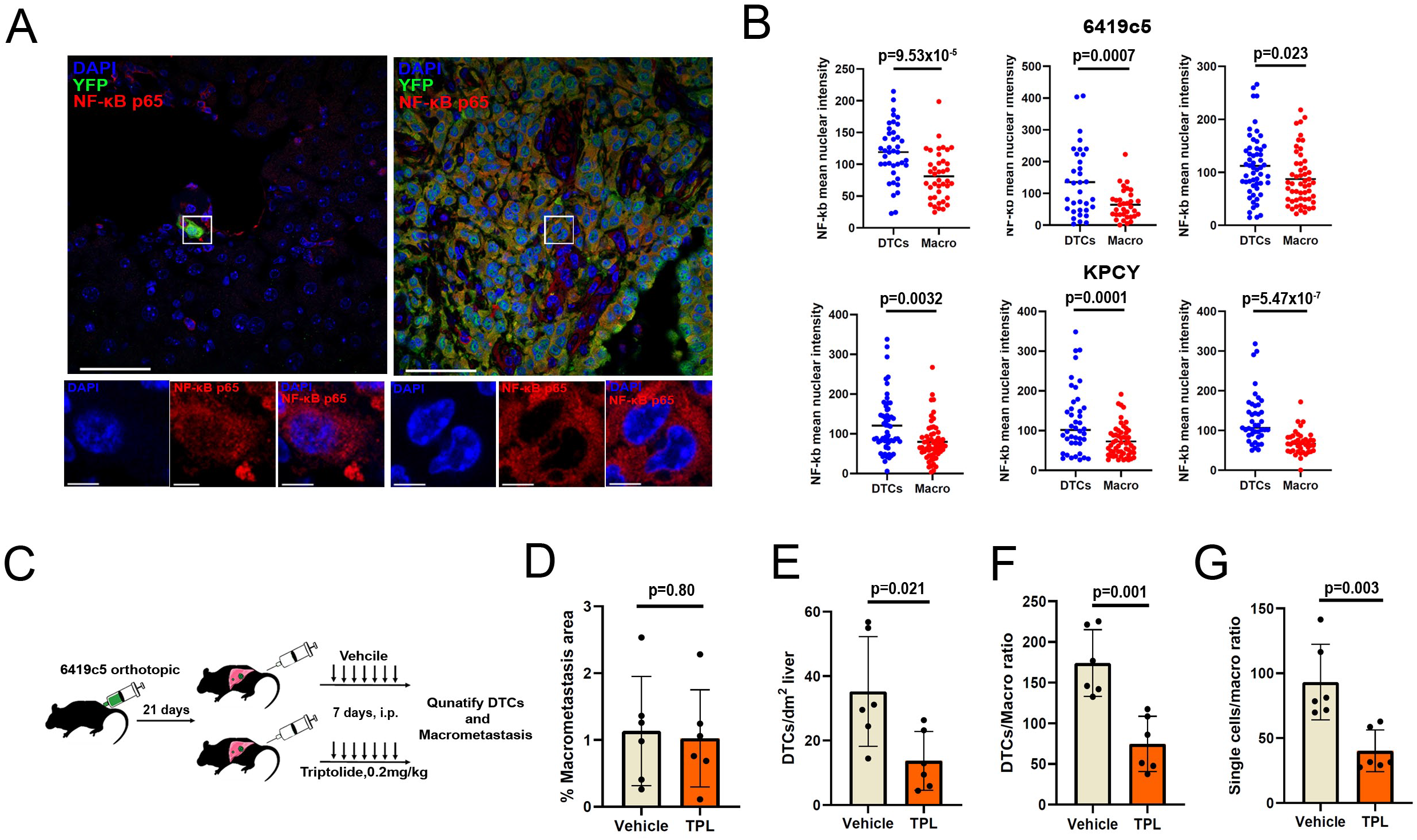
DTCs are susceptible to inhibition of NF-κB /AP-1 signaling. **(A)** Representative confocal images of DTCs and macrometastasis from livers of KPCY mice stained with NF-κB (red) and YFP (green) antibodies. Scale Bar=50 μm. On the bottom, higher magnification of inset showing nuclear signal of NF-κB in 1μm z-section. NF-κB (red) and DAPI (blue) Scale Bar=5 μm. **(B)** Quantification of mean nuclear intensity of NF-κB in DTCs and macrometastasis of orthotopic 6419c5-YFP (top) and KPCY mice (Bottom). Each plot represents a single animal **(C)** Experimental scheme for testing NF-κB /AP-1 targeting by Triptolide (TPL) in neo-adjuvant settings. **(D-G)** Quantification of metastatic colonies in livers of TPL and vehicle treated mice. **(D)** Percentage macrometastasis area of liver area **(E)** Absolute DTCs frequency in liver sections. **(F)** Ratio between DTCs frequency and area of macrometastasis. **(G)** Ratio between single-cell DTCs frequency and area of macrometastasis (n=6 animals per group, 10 liver sections were sampled for each animal). Bars represent mean ± SEM in all graphs. p-values were calculated by unpaired two-tailed Student’s t-test.

## Discussion

Studying the occult fraction of metastatic disease is challenging due to the rarity of cells, and lack of efficient tools to isolate and define them. PIC-IT overcomes these challenges, allowing us to isolate and systematically characterize the function of tumor cells derived from the earliest stages of spontaneously arising metastasis. Our experiments reveal that DTCs are a diverse cell population enriched with cells manifesting a pro-inflammatory response. Our findings further highlight a specific role for NF-κB/AP-1-regulated inflammatory pathways in DTCs not shared by the majority of metastatic cells (macrometastasis), demonstrating the utility of our approach in identifying susceptibilities of a micro-fraction of the metastatic population that drives disease recurrence and mortality. Moreover, the live cell isolation feature of PIC-IT allows a direct measurement of the proliferative capacities of these two cell populations, suggesting that the presence or absence of a “poised” proliferative state may contribute to a metastatic lesion’s propensity to grow rapidly or slowly (or to become dormant)(Maria Soledad Sosa, Bragado, Debnath, & Aguirre-Ghiso, 2013).

While the current study used PIC-IT to isolate cells at different stages of metastasis, the method has many potential applications. For example, it could be used to isolate cells in different “niches” of the tumor microenvironment (e.g. cells in well perfused vs. poorly perfused regions), thus paving the way for testing functional features such as proliferation, migration, metabolic characteristics, and drug resistance in heterogeneous tumor cell populations. As our identification of CLCA1 shows, the technique can be used to identify protein markers – particularly those residing on the cell surface – with which to prospectively isolate cells of interest by flow cytometry. Finally, the technique is particularly well suited to the isolation of spatially-defined rare cells – such as stem cells – in malignant and benign conditions.

## Materials and methods

### Cell culture

Pancreatic tumor cells were isolated from late-stage primary tumors from C57BL/6 background KPCY mice and generated by limiting dilution as described(Li et al., 2018). All mouse pancreatic tumor cell clones were tested by the Research Animal Diagnostic Laboratory (RADIL) at the University of Missouri, using the Infectious Microbe PCR Amplification Test (IMPACT) II. Tumor cells were cultured in DMEM (high glucose without sodium pyruvate) with 10% FBS (Gibco) and glutamine (2mM). The clones were regularly tested using the MycoAlert Mycoplasma Detection Kit (Lonza).

### Genetic modification of tumor cells

6419c5-YFP cells were transduced with a sgRNA targeting endogenous YFP to remove its expression (sgRNA sequence: GGGCGAGGAGCTGTTCACCG). Subsequentially, 6419c5 and 5074 cells were transduced with a lentiviral expression vector of h2b-dendra2 (Addgene #51005). Single cell clones expressing Dendra2-H2B were derived by FACS sorting.

### Animals

Wild-type C57BL/6 were purchased from The Jackson Laboratory and/or bred at the University of Pennsylvania. For tumor-bearing mice, endpoint criteria included severe cachexia, weakness and inactivity.

### Orthotopic Implantation of tumor cells

Tumor cells were dissociated into single cells with 0.25% trypsin (Gibco), washed with serum-free Dulbecco’s Modified Eagle’s medium (DMEM) twice, and counted in preparation for orthotopic implantations. 3,000-10,000 tumor cells were implanted orthotopically into 6-8-week old female or male C57BL/6 mice. Tumors were harvested 3-5 weeks following implantation.

### Triptolide treatment

10,000 6419c5-YFP tumor cells were implanted orthotopically and tumors were allowed to grow for 3 weeks. Mice were randomized into two groups and injected intraperiotenaly with either 0.2mg/kg Triptolide (HY-32735, Medchem express) or Vehicle (10% DMSO, 40% PEG300, 5% Tween-80, 45% Saline) for 7 consecutive days. Livers were sectioned and DTC frequency represent 10 sections per mouse. Macrometastatic area and total liver area were measured using imageJ.

### PIC-IT

Tumor bearing mice were sacrificed and livers or tumors were removed and fragmented using scissors to small pieces (~2-5 mm in diameter) and kept in flow buffer (HBSS) on ice. Fragments were further flattened and mounted on top of a petri-dish, covered with a drop of flow buffer (HBSS,2.5mM HEPES 5% FBS, 2mM glutamine, 1% Penicillin/streptomycin, 1% Nonessential amino acids, 0.5% Fungizone, 0.3% Glucose, 20mM Mgcl2, 1mM Sodium pyruvate). and examined under a BX60 Olympus upright microscope using a x10 objective unless otherwise specified. Metastasis were identified and FOV was confined around metastatic foci of choice using the field stop element. Cells were then photoconverted by illumination with a wavelength of 400-450nm using mercury lamp and a 451/106 nm BrightLine^®^ Full Spectrum Blocking single-band bandpass filter (FF01-451/106-25, Avro Inc) and red fluorescence was confirmed. Photoconverted specimen of each metastatic group was pooled and transferred to a C-column (130-093-237, Miltentyi Biotech) and digested by incubation with 2mg/ml collagenase IV+DNase I (Sigma) in DMEM using the mLIDK program of a gentleMACS (Miltentyi Biotech). Cells were mashed on 100μm MESH strainer, pelleted and dissociated to single cells using

Accutase (07922, Stemcell technologies) for 5’, countered, pelleted an incubated for 3’ with ACK lysis buffer(10-548E, Lonza) to remove red blood cells. Pelleted cells were resuspended with Flow buffer containing 1ug/ml DAPI +DNase I, incubated for 5’ in 37°C, filtered through 100μm mesh strainer and either analyzed by LSRII or sorted using a FACSaria. Isolation experiments represent independent experiments performed on different days.

### Spatial analysis

For spatial analysis, primary tumors were processed through a VT1200S vibratome (Leica Biosystems) to 200 μm slices. One maximally confined FOV of X10 objective was photoconverted per slice and Z-stacks were acquired by a LEICA TCS SP8 microscope. Sum intensity projections from each sample were analyzed by ImageJ using the radial profile plugin.

### Flow cytometry and cell sorting

For flow cytometric analyses, dissociated livers were stained with Armenian hamster anti-CLCA1 1:5 (10.1.1, Iowa Hybridoma bank) for 25’ following ACK lysis. Samples were then washed and stained with an APC-conjugated secondary antibody and PE-Cy7-CD45 antibody (103114, Biolegend) and then analyzed by flow cytometry using BD FACS LSR-II (BD Biosciences) and analyzed FlowJo software (Treestar).

For cell sorting, digested organs were sorted into 96wells prepped for cel-seq2 or 96 well culture dishes for re-culturing experiments using FACSariaII equipped with a green laser through a 100μm nozzle. Isolation experiments represent 3 experiments performed on different days.

### Cel-seq2 profiling

Library preparation was performed by adhering to the developer’s protocol using the original barcodes described (Hashimshony, 2016 genome biology). Libraries were quantified using qubit (Thermo fisher) and library fragmentation was assessed via tapestation (Agilent). Libraries were prepared for sequencing using Illumna 75 cycles High output kit v2.0 (20024096, Illumna) and sequenced on a NextSeq 500/550 with 26 bases on read1 (R1), 8 bases for the Illumna index and 41 bases for read 2 (R2).

### Immunofluorescent and immunohistochemistry staining

For immunostaining, tissues were fixed in Zn-formalin for 24 hours and embedded in paraffin. Sections were deparaffinized, rehydrated and prepared by antigen retrieval for 6 minutes each, and then blocked in 5% donkey serum for 1 hour at room temperature, incubated with either Rabbit anti-BNIP3 antibody at 1:100 (A5683, Abclonal), Rabbit anti-NF-kB-p65 1:400 (8242S, Cell signaling technology), Rabbit anti-Phospho-c-jun 1:200, (3270S, Cell signaling technology), Rat anti-ki67 1:100 (14-5698-82 ebiosciences) and Goat anti-GFP (ab6673, Abcam) at 1:250 overnight at 4°C, washed, incubated with secondary antibody AF594-anti-rabbit (A-21207, Invitrogen), Alexa594 anti-rat (A-21209, Invitrogen) and Alexa488 anti-goat (11055, Invitrogen) antibodies at 1:200 for 1 hour at room temperature, counterstained with DAPI, washed and mounted. Slides were visualized and imaged using an Olympus IX71 inverted multicolor fluorescent microscope and a DP71 camera using via x400 magnification. MFI was calculated by measuring average intensity over a fixed threshold for all images. NF-kB staining was imaged via Zeiss LSM880 confocal microscope using X63 oil-immersion objective and intensity was measured on 0.8μm z-sections.

### Computational analysis

Fastq files were generated from bcl files using bcl2fastq tool from illumina. Reads were then trimmed to 12 bp for forward sequencing (R1) and 36 bp for reverse sequencing (R2), using cutadapt software (https://cutadapt.readthedocs.io/en/stable/). Trimmed reads were first debarcoded using UMI_tools with “extract” function and default settings (https://github.com/CGATOxford/UMI-tools)and then aligned to mm10 genome using STAR (https://github.com/alexdobin/STAR). Aligned reads were counted using featureCounts (http://subread.sourceforge.net) and output as bam files. Generated bam files were then sorted and indexed using samtools. Molecules were counted using UMI_tools with “count” function and default settings to generate UMI tables. The merged UMI table was subsequently imported into R-studio (R version 3.3.3) and used as input for edgeR analysis to assess differentially expressed genes. Samples with 20,000–60,000 UMI were used for differential gene expression analysis. Differentially expressed genes were also used as input file for online GSEA analysis following provided instruction. Detailed Scripts and parameters used for each steps of analysis could be provided by reasonable request to the authors. Gene signatures were extracted by performing edgeR analysis.

For visualization of the single cell RNA-seq results, we used the Seurat package. We performed SCT transformation for data normalization and integration. We then performed linear dimensional reduction using PCA, followed by clustering using K-nearest-neighbor graph-based clustering and Louvain modularity optimization (with dimensions= 1:15 and resolution=1), and non-linear dimensional reduction using UMAP method. Markers for each identified clusters were generated using Seurat with default settings and used as input for function pathway analysis, including online GSEA analysis using the molecular signatures database.

For motif analysis: Top differentially expressed genes were used as input for HOMER (−1000bp to +300bp as scanning region) for identification of motifs of potential transcriptional regulators. RNA-seq and clinical data of human pancreatic cancer samples from TCGA dataset were utilized for survival analysis. PDAC samples were first scored for enrichment of indicated gene signatures using GSVA. Clinical information associated with top and bottom 25% patients based on the GSVA score of each gene signature were visualized as survival plots using Prism.

### Ex vivo culture assay and colony formation

Sorted tumor cells were seeded at a density of 45 cells per 96/well on Growth factor-reduced Matrigel (BD) and allowed to grow for up to 6 days under standard culture conditions (See cell culture). Retention was estimated 24hrs post seeding. For colony size quantifications, cultures were evaluated daily and the number of cells was manually counted per colony. Isolation experiments represent 3 experiments performed on different days.

### Software and Statistical analysis

PRISM software and R were used for data processing, statistical analysis, and result visualization (http://www.graphpad.com). The R language and environment for graphics (https://www.r-project.org) was used in this study for the bioinformatics analysis RNA-seq. The R packages used for all analysis described in this manuscript were from the Bioconductor and CRAN. Statistical comparisons between two groups were performed using Student’s unpaired t test. For comparisons between multiple groups, one-way ANOVA with Tukey’s HSD post-test was used. If outliers were indicated by the software, they were removed. Otherwise, all relevant data was included. On graphs, bars represent either range or standard error of mean (SEM), as indicated in legends. For all figures, p<0.05 was considered statistically significant, * indicates p<0.05, **p<0.01, and ***p<0.001.

