## Supplementary Material for "Dissecting phenotypic transitions in metastatic disease via photoconversion-based isolation"

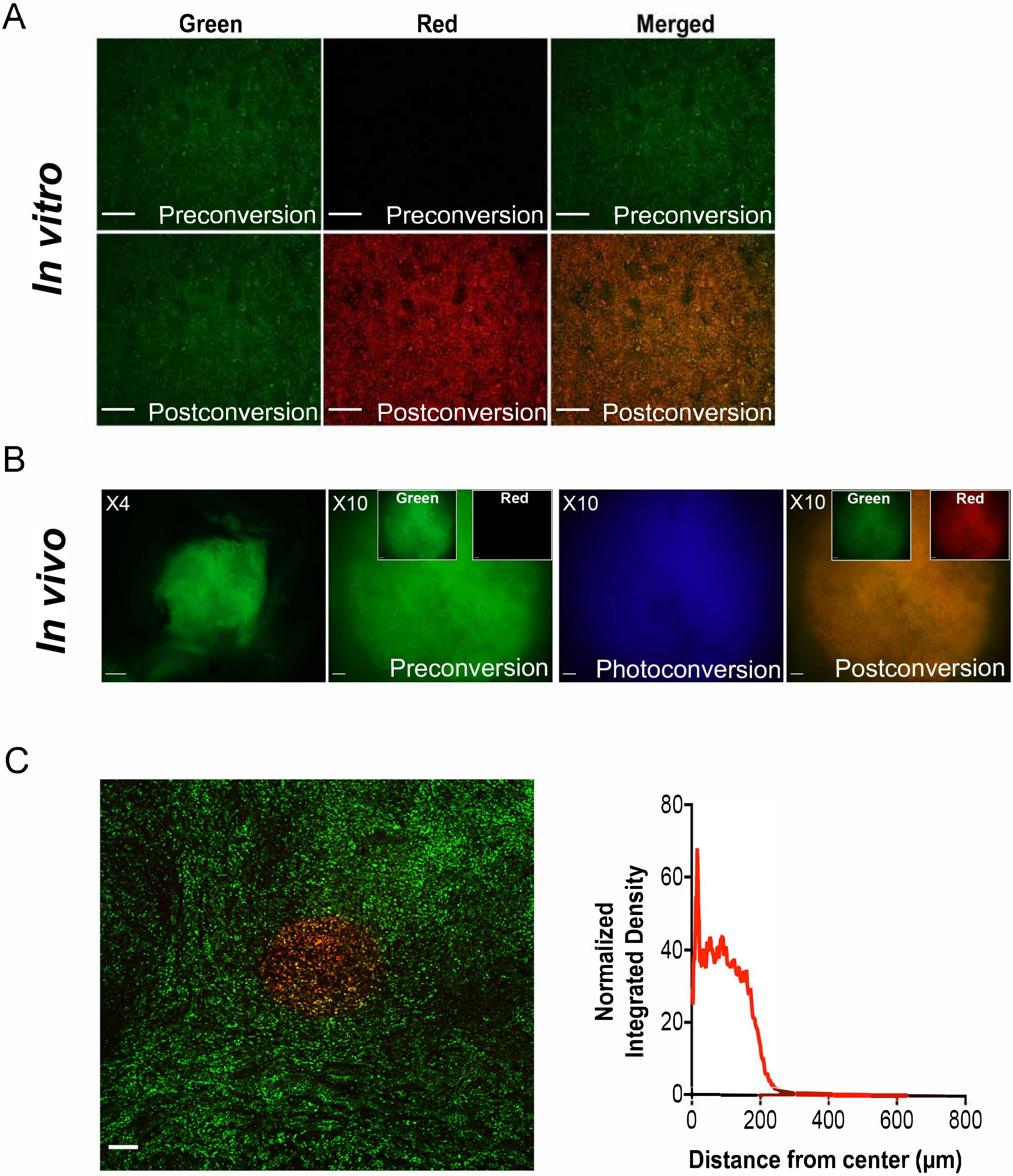


Figure1-figure supplement 1

**Figure1-figure supplement 1. Spatial flexibility of PIC-IT**

**(A)** Photoconversion of cultured PDA tumor cells. Representative images of pancreatic tumor cells preconversion and following 30 seconds exposure mercury lamp-generated violet light *in vitro* (10X objective, Scale Bar = 100 µm). **(B)** Photoconversion of Dendra2 in spontaneously arising macrometastasis. Left to right: Low magnification microscopic view of a Dendra2-positive macrometastasis before conversion (4X objective, Scale Bar=250µm). FOV of the lesion prior to photoconversion (10X objective, Scale Bar=100µm). In boxes, images representing individual green and red channels for the field-stop FOV. Image acquired during photoconversion session. FOV of the macrometastasis following photoconversion. **(C)** Spatial resolution of photoconversion. Representative confocal microscopy image of a precision slice from a liver metastasis photoconverted in one maximally confined FOV using a 10X objective (left panel, Scale bar = 100µm) and radial intensity profile derived (n=10) for Dendra2-red channel of the photoconverted FOV (right panel).

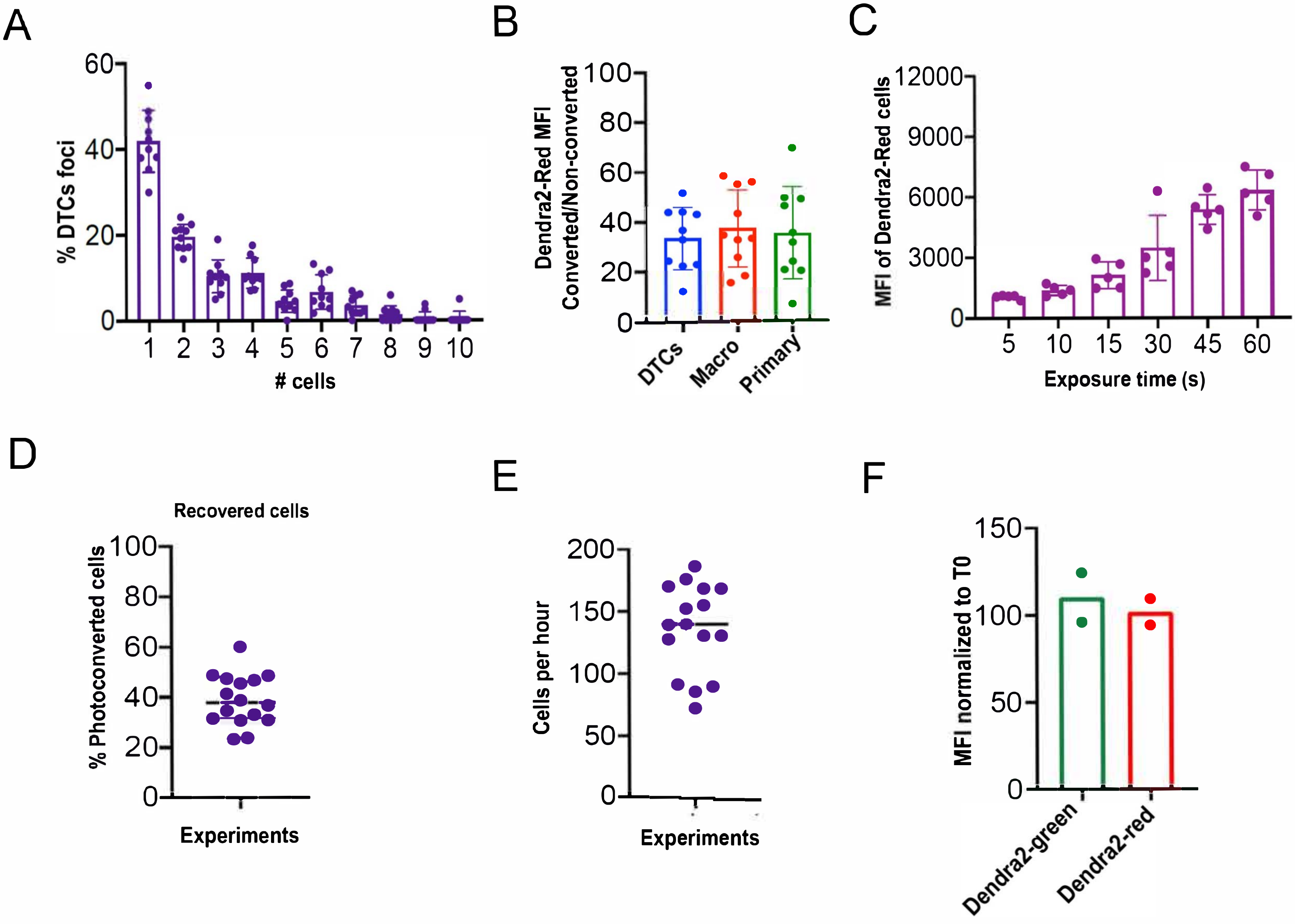


Figure1-figure supplement 2

**Figure1-figure supplement 2. Kinetics of cell marking and efficiency of isolation using PIC-IT.**

**(A)** Size distribution of DTCs collected in isolation sessions. Metastatic foci of 1-10 cells were binned according to size and the frequency of each size plotted as a percentage of the total across 10 independent experiments. **(B)** Average fold-change in red signal over background of non-converted samples in metastatic and primary tumors (n=10 independent experiments). **(C)** Fold-change increase in Dendra2-red in photoconverted primary tumor cells as a function of time of exposure (seconds) to violet light. Data represents an average of 3 technical replicates per timepoint. **(D)** Recovery of tumor cells from DTC foci calculated as the ratio of the number of photoconverted cells recovered by flow cytometry to the number of cells photoconverted in each microscopy session (n=16 sessions). **(E)** Average number of tumor cells recovered from DTCs per hour photoconversion session. (n=16). **(F)** Stability of Dendra2-red in photoconverted metastatic tumor cells. Ratios of Dendra2-green and Dendra2-red intensities between macrometastasis-derived photoconverted cells processed and analyzed by flow cytometry 7 hours and 5 minutes post conversion. Ratio of fractions of Dendra2-red cells at 7 hours and 5 minutes (n=2 independent experiments **(**Bars represent mean+SEM in all graphs .

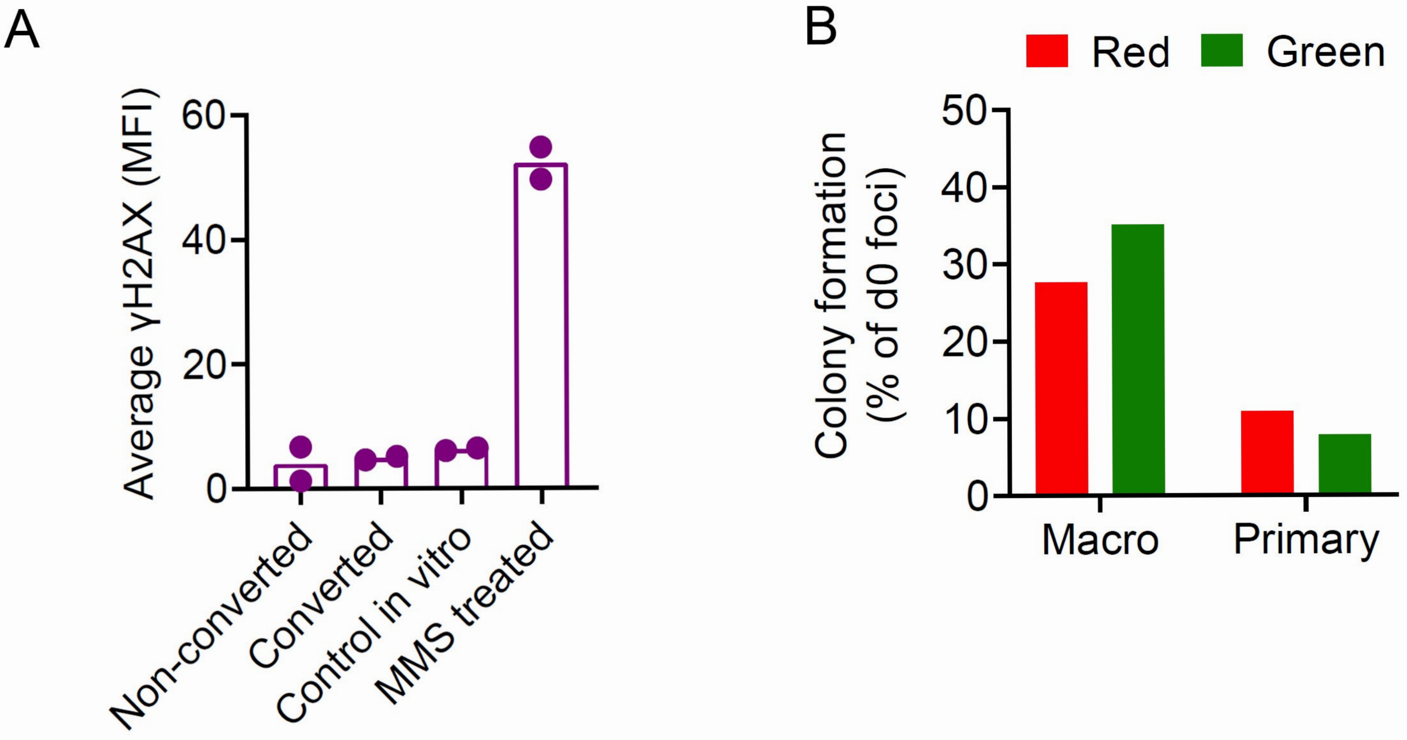


Figure1-figure supplement 3

**Figure1-figure supplement 3. Viable isolation of tumor cells by PIC-IT.**

**(A)** Evaluation of DNA integrity of isolated metastatic cells. Immunostaining for γ-H2AX staining intensity (MFI, Mean fluorescence intensity) in tumor cells at 24hrs post isolation (n=2). **(B)** Comparison of colony formation *ex vivo* of photoconverted and non-converted cells derived from macrometastasis or primary tumors 6 days post-seeding (n=2 independent experiments and 6 technical replicates for each group). A colony was defined as a cell cluster >10 cells.

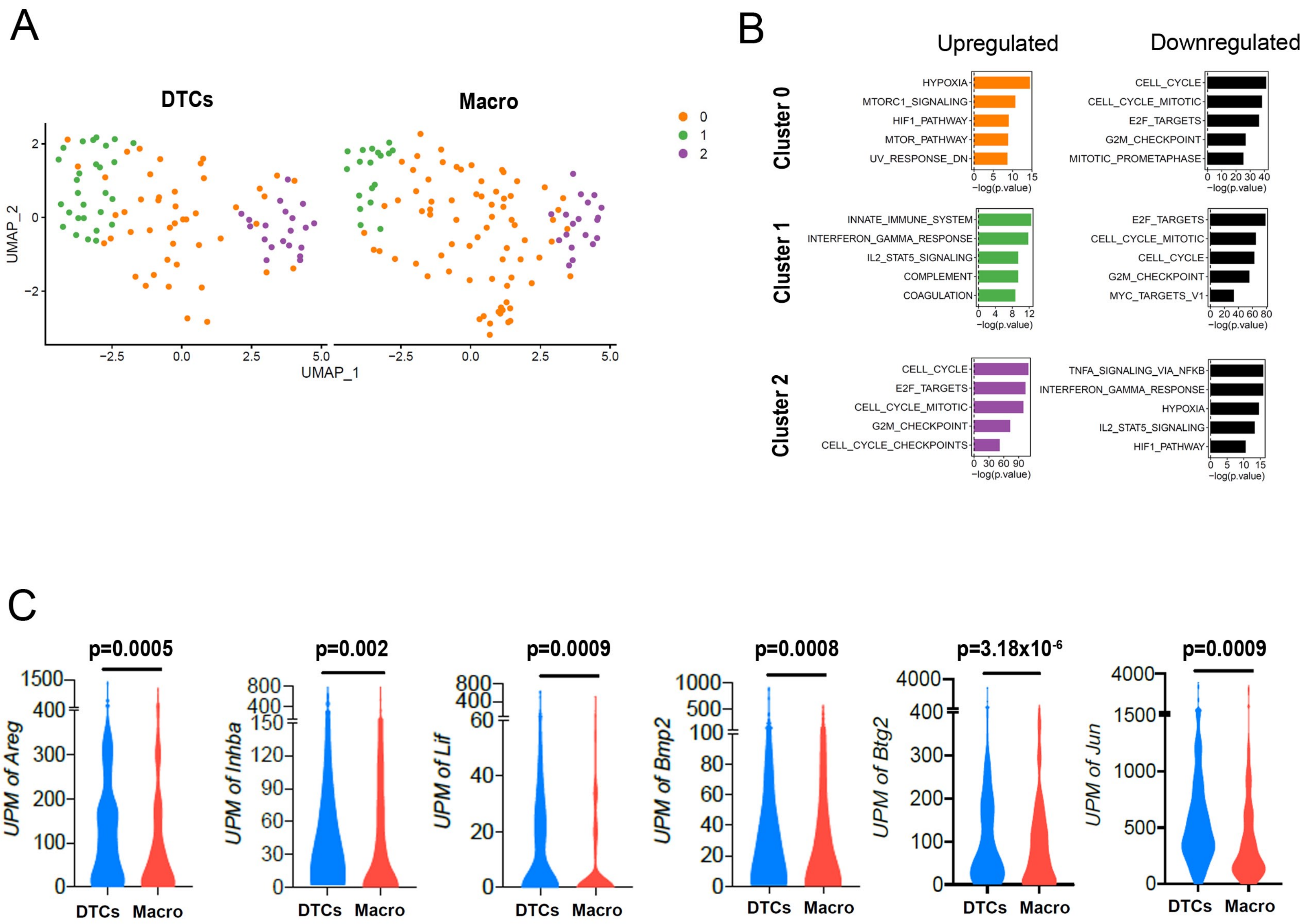


Figure2-figure supplement 1

**Figure2-figure supplement 1. Phenotypic transition from DTCs to macrometastasis. (A)** Individual view of DTC and macrometastasis clustering visualized by UMAP. **(B)** Functional annotation for Hallmark and canonical pathways via GSEA msGDMIB ($p<0.05$, 200 highest fold-change genes in each cluster). **(C)** Abundance of transcripts in RNA-seq for transcriptional targets of NF-κB significantly upregulated in DTCs. ($N_{DTCs}=94$, $N_{macro}=110$). p-values were calculated by unpaired two-tailed Student's t-test.

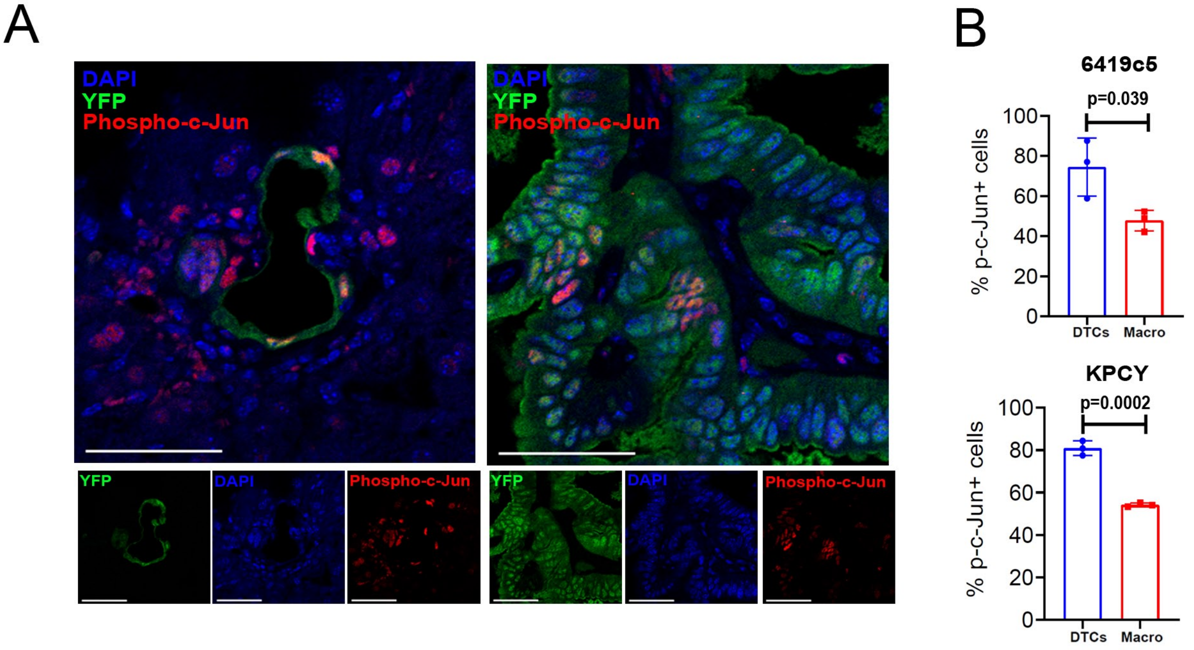


Figure 4-figure supplement 1

**Figure4-Figure supplement 1. Increased activity of c-JUN in DTCs.**

**(A)** Representative confocal images of DTCs and macrometastases from livers of KPCY mice stained with phosphorylated c-Jun (red) and YFP (green) antibodies. Scale Bar=50 μm. On the bottom, individual channels. **(B)** Fraction of Ser73-phosphorylated c-jun positive cells in DTCs and macrometastasis of orthotopic 6419c5-YFP (top) and KPCY mice (Bottom). (n=3 animals per group). Bars represent mean ± SEM in all graphs. p-values were calculated by unpaired two-tailed Student's t-test.
